# Sporodochia formed by *Fusarium oxysporum* f. sp. *fragariae* produce airborne conidia and are ubiquitous on diseased strawberry plants in California

**DOI:** 10.1101/2022.10.13.512161

**Authors:** Peter M. Henry, Christine J. Dilla-Ermita, Polly Goldman, Jose Jaime, Gerardo Ramos

## Abstract

Sporodochia are dense masses of fungal hyphae bearing asexual conidia. For *Fusarium oxysporum*, sporodochia are known to produce airborne conidia and enhance the dissemination of this otherwise soilborne pathogen. Sporodochia are small and transient, and they are documented for only a few *formae speciales* of *Fusarium oxysporum*. This study reports airborne conidia and sporodochia produced by *F. oxysporum* f. sp. *fragariae*, the cause of Fusarium wilt of strawberry, in the Monterey Bay region of California. Sporodochia were consistently present in Fusarium wilt-afflicted strawberry fields and were discovered in 21 of 24 Fusarium wilt-diseased fields. Only necrotic tissues were observed bearing sporodochia, and they were most frequently observed on petioles and peduncles. Sporodochia grew significantly longer up peduncles than petioles, extending further away from the base of the plant and toward parts of the canopy more exposed to wind. A stolon hosted the longest stretch of sporodochial growth, found covering the stolon’s entire 35 cm length and also the base of the daughter plant. Macroconidia were produced by all sporodochia samples, and we did not find microconidia on any samples. An initial series of experiments confirmed the potential for conidia produced by sporodochia to disperse with wind over short distances. The prevalence of sporodochia producing airborne spores of *F. oxysporum* f. sp. *fragariae* has great importance for disease management and biosecurity.

---

Fusarium wilt of strawberry is a widespread disease that causes severe yield losses. In California, where 90% of the United States’ fresh market strawberries are grown, this disease is caused by the highly virulent ‘yellows-*fragariae’* pathotype of *Fusarium oxysporum* f. sp. *fragariae* (Henry et al. 2017; Henry et al. 2021). This disease was first reported in coastal California in 2006 and has since become widespread, particularly in the Monterey Bay growing region where Fusarium wilt-susceptible cultivars are common (Koike and Gordon 2015; Koike et al. 2009).

*Fusarium oxysporum* is typically considered a soilborne fungus that disperses with soil and water or on infested plant material. However, *F. oxysporum* f. sp. *lycopersici* (tomato wilt), *basilici* (basil), *radicis-lycopersici* (tomato crown and root rot), and *radicis-cucumerinum* (cucurbit crown and root rot) can produce airborne conidia from sporodochia formed on the surface of infected plant tissues (Katan et al. 1997; Rekah et al. 2000). Airborne conidia can facilitate dispersal to new locations and rapidly colonize steam pasteurized soil (Rowe et al. 1977). For *F. oxysporum* f. sp. *radicis-lycopersici* and *basilicum*, the airborne conidia can also initiate foliar plant infections leading to systemic plant colonization (Rekah et al. 2000). Sporodochia may attract insects, such as fungus gnats (e.g. *Sciaria* spp.), that can acquire spores and vector them to new locations (Scarlett et al. 2014). To our knowledge, there have been no previous reports of sporodochia produced by *F. oxysporum* f. sp. *fragariae*.

On Aug 4, 2022, we observed orange and cream-colored fungal masses on necrotic strawberry plants at a field with high mortality and a history of Fusarium wilt. These fungal masses were found on the necrotic tissues of almost all wilting plants in the field. Upon microscopic observation, these masses yielded abundant macroconidia and single-spored colonies tested positive with qPCR for the yellows-*fragariae* pathotype of *F. oxysporum* f. sp. *fragariae*. Other pathogens causing strawberry wilt (*Macrophomina phaseolina, Verticillium dahliae*, or *Phytophthora* spp.) were not present in crown extracts tested with recombinase polymerase amplification (RPA); the same procedure yielded a positive result for *F. oxysporum* f. sp. *fragariae*. We concluded these structures were sporodochia produced by *F. oxysporum* f. sp. *fragariae* on necrotic, above-ground strawberry tissues.

Subsequently, we initiated a project to explore the significance of this discovery by: 1) documenting the prevalence of sporodochia in Fusarium wilt-afflicted fields in the Monterey Bay strawberry production region, and 2) conducting a preliminary evaluation of the potential for aerial dispersal of *F. oxysporum* f. sp. *fragariae* spores across short distances. Here, we show that sporodochia are ubiquitous on Fusarium wilt-afflicted plants across fields in the Monterey Bay region of California, and airborne dispersal could partially account for the prevalence of the disease in this area.

After the initial observation, a total of 34 strawberry fields with wilt disease were visited between August 4, 2022 and September 26, 2022. Symptomatic plants were examined in-field and without magnification for sporodochia. If fungal growth similar in morphology to sporodochia was observed, tissues bearing these structures were saved for laboratory analysis. From each field we also collected the crowns of eight symptomatic plants for recombinase polymerase amplification (RPA) assays.

For plants observed to have sporodochia, we recorded: the type of tissue from which the sporodochia grew, the length of sporodochia production along the sample, and whether micro-or macroconidia were recovered from these structures. Differences between tissue types in the length of sporodochia were statistically tested in R (version 3.6.1) with a Kruskal-Wallis Rank Sum test (package=‘kruskal.test’) and pairwise Wilcoxon Rank Sum Tests with Benjamini Hochberg correction for multiple comparisons package=‘pairwise.wilcox.test(p.adjust.method=“BH”) (R_Core_Team 2013). Microscopic analysis was conducted by gently scraping sporodochia with a sterile scalpel and staining the conidia with lactophenol cotton blue for at least 15 minutes. Conidia were examined under a compound microscope with 100 to 400 × magnification. Additionally, we prepared a composite spore suspension (two plants per field sample) for single-spore culturing. To accomplish this, a sterile surgical blade was used to gently scrape the surface of the single sporodochium from each plant and transfer to a microcentrifuge tube with 600 µl of sterile, de-ionized water. This solution was diluted ten-fold and 100 µl spread evenly across a 100 mm plate of Komada’s medium (Komada 1975). For each sample, three single colonies emerging from the Komada’s medium after seven days were selected for molecular identification of *F. oxysporum* f. sp. *fragariae*. DNA was extracted from each colony using the Prepman Ultra (ThermoFisher Scientific) reagent following the manufacturer’s recommended protocol. The qPCR-based identification protocol published by Burkhardt et al. (2019) was used to determine if the single-spored colony was *F. oxysporum* f. sp. *fragariae*.

The 8-crown composite sample was used to determine if *F. oxysporum* f. sp. *fragariae* was a causal agent on plants from which sporodochia samples were obtained. For this procedure, 1) a total of 1g of discolored crown tissues were collected from the eight plants, 2) a crude DNA solution from this 1g sample was extracted in 10 mL GEB2 buffer by the method described in Burkhardt et al. (2019), 3) a single replicate of a recombinase polymerase amplification assay was conducted for each of the following strawberry wilt pathogens: *F. oxysporum* f. sp. *fragariae, Macrophomina phaseolina, Verticillium dahliae*, and *Phytophthora* spp. (Bilodeau et al. 2012; Burkhardt et al. 2019; Burkhardt et al. 2018; Miles et al. 2015). Two 1 g samples were collected for each field; therefore, each field was twice assayed for each of the four pathogens. All RPA assays were run with an internal control reaction targeting the *COX* gene with universal plant-specific primers. If both the control and target reaction did not amplify, the assay was re-run.

We conducted a simple experiment to evaluate the potential for conidia produced on sporodochia to be aerially dispersed. A “spore dispersal box” was prepared by creating a 15.7 cm hole in both sides of a Sterilite 25-quart plastic box. A 15.7 cm fan was placed in one hole and a Hoover Type Y allergen filtration bag was secured to the hole on the other side. The box was cleaned with 70% ethanol before the start of each trial. Fresh sporodochia samples were collected and incubated in a high humidity chamber under continuous fluorescent lighting for two days to enhance sporulation. Four experiments were conducted, each using a different sporulating plant sample. The spore dispersal box was placed in a controlled environment chamber with continuous lighting and a 30° temperature setting. Inside the box, a plant sample was placed between the fan and ten labeled plates of Komada’s medium that had been moistened evenly with 200 µl sterile de-ionized water (Schweigkofler et al. 2004). Six plates were arranged on the bottom of the box while four plates were affixed to the sides. The lid of the box was partially closed and the fan was run at maximum speed, generating wind speeds of 7.56 to 8.85 kilometers per hour for 120 minutes. Plates were then incubated at 22.8°C under continuous lighting for seven days. Colonies were then counted and five representative colonies per experiment were diagnosed as *F. oxysporum* f. sp. *fragariae* by DNA extraction and qPCR as described above.

Sporodochia were observed on plants from 21 of the 34 fields; *F. oxysporum* f. sp. *fragariae* was diagnosed as a causal agent for wilt disease at each of these 21 fields (Table 1). Almost all plants with necrotic petiole or peduncle tissues bore sporodochia at the 21 fields. The sporodochia were observed on four Fusarium wilt-susceptible cultivars planted: ‘Monterey’, ‘Cabrillo’, ‘Albion’, and ‘Sweet Ann’. Sporodochia were not observed in any field that tested negative for *F. oxysporum* f. sp. *fragariae* in crown RPA assays (data not shown). There were only three fields where sporodochia were not observed but the crown RPA assay was positive for *F. oxysporum* f. sp. *fragariae* (Table 1). For two of these fields, the RPA assay was also positive for *Macrophomina phaseolina*, and one of the fields was planted with a Fusarium wilt-resistant cultivar, ‘San Andreas’ (Table 1) (Pincot et al. 2018).

**Table 1.**
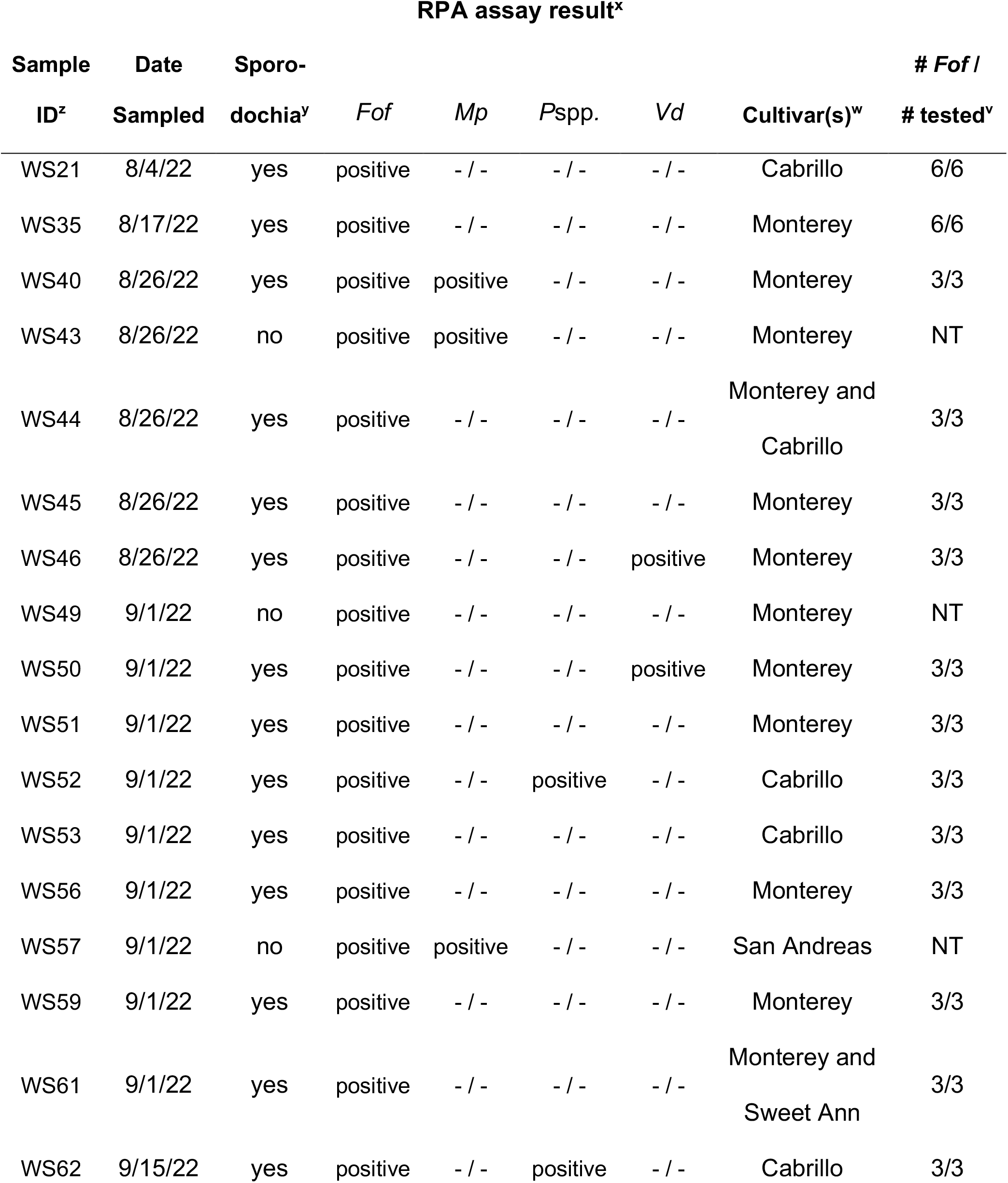
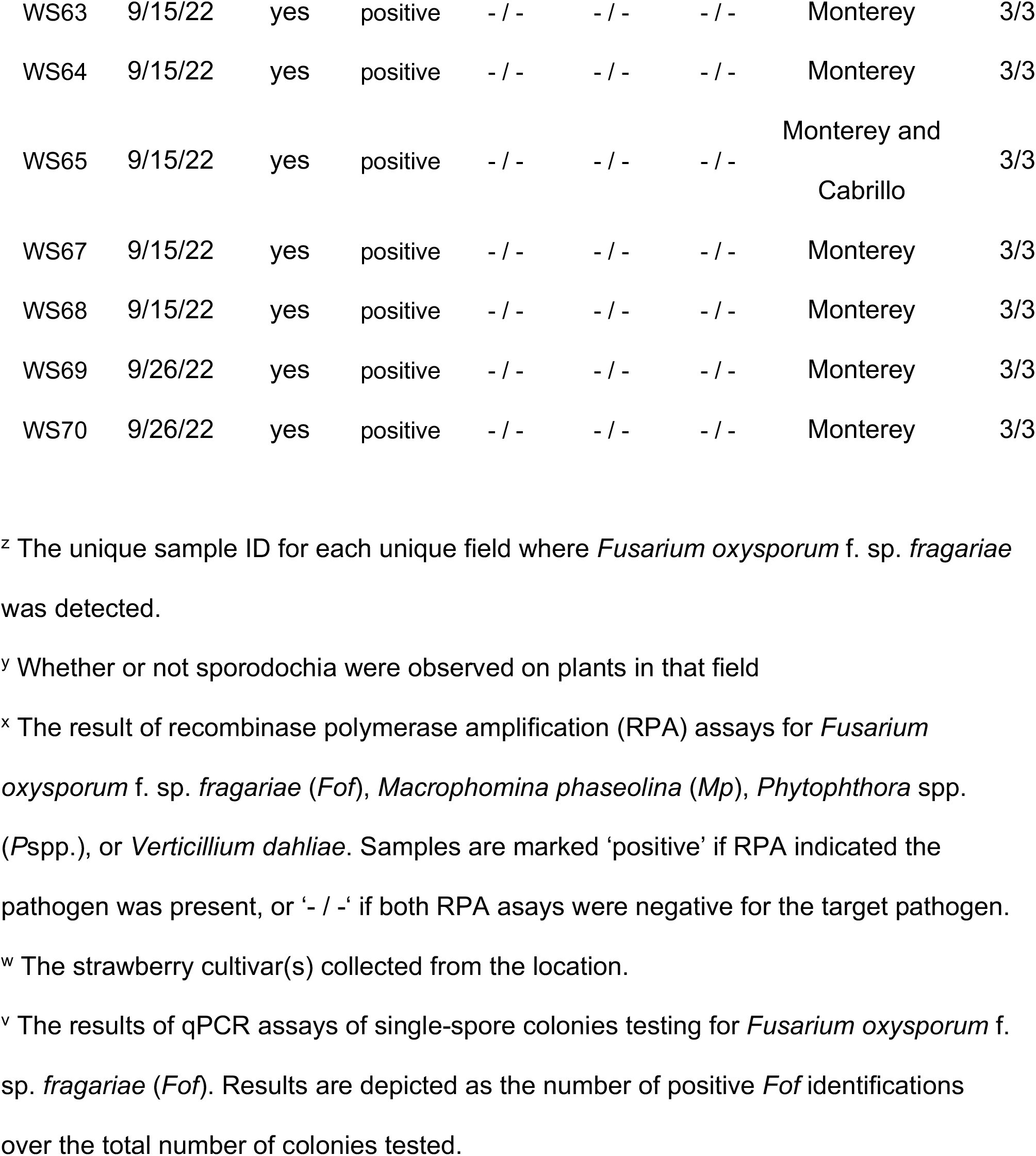
Summary of results from sporodochia and crown samples.

Necrotic petioles, peduncles, and stolons all bore sporodochia; this structure was never observed on living tissues (Figure 1; Table 1). Petioles were the most common site of sporodochia, especially at the base of the petiole where it is attached to the crown and fully encased by plant tissues. Although the sporodochia occasionally appeared to grow from the crown, closer inspection revealed they emerged from the outside of petiole bases and not the crown cortex. Significantly greater lengths of sporodochia were observed covering peduncle samples than petioles or petiole bases (*P* ≤ 0.009; Figure 1; Table 1). The maximum length of sporodochia along peduncle tissues was 15 cm. The only stolon sample collected in this study had 35 cm of sporodochia that extended from the mother crown along the stolon and onto the base of the daughter plant (Figure 1C). Macroconidia were recovered from all sporodochia, and microconidia produced by *F. oxysporum* f. sp. *fragariae* were not observed (Figure 2). Single-spored colonies from each sporodochia sample were confirmed to be *F. oxysporum* f. sp. *fragariae* by qPCR (Table 1).

**Fig. 1.**
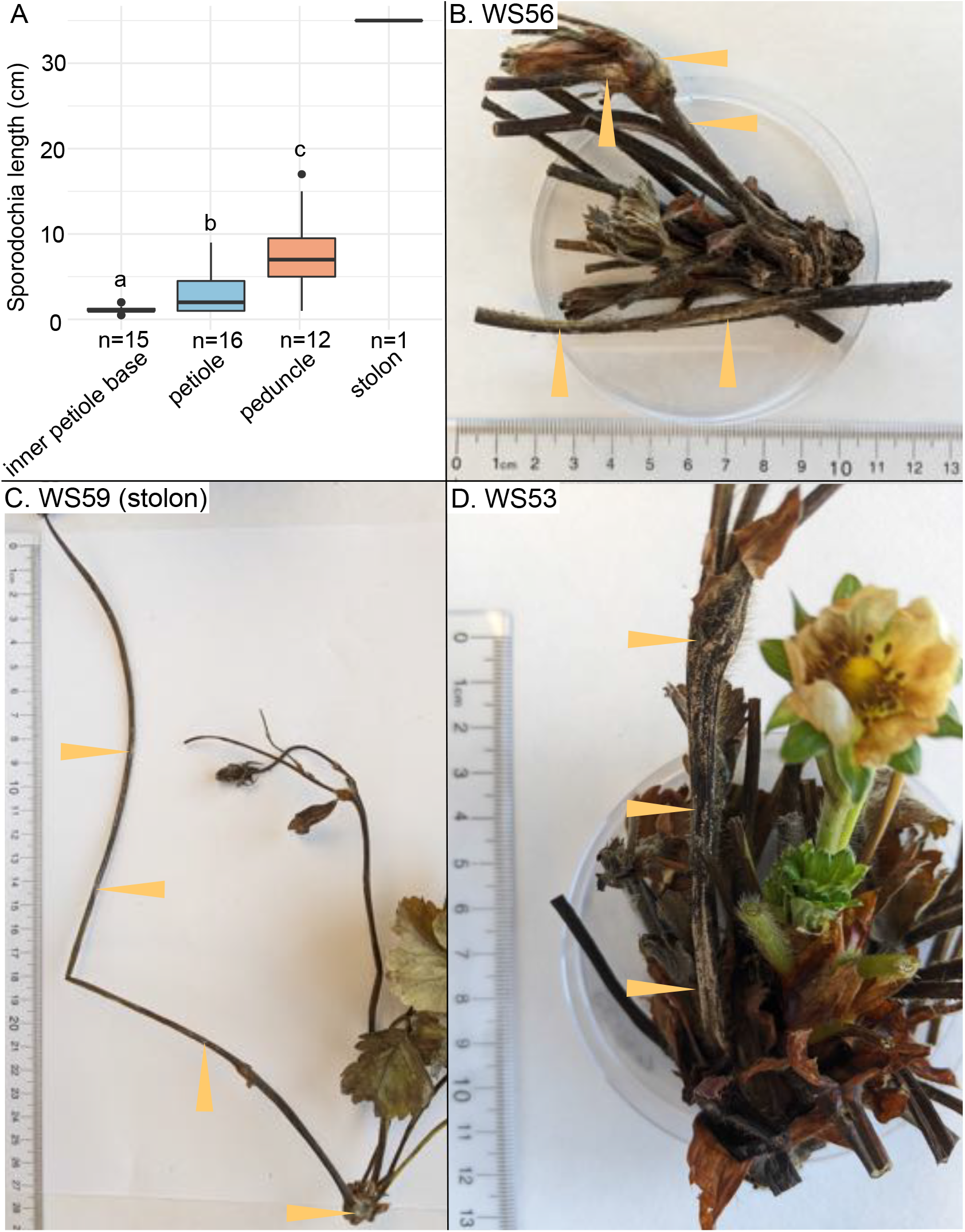
Macroscopic description of strawberry tissues bearing sporodochia. **A** – a box-plot depicting the length of sporodochial growth along strawberry tissues. The number of samples depicted is indicated with ‘n=#’ above the x-axis label. Different letters indicate significant differences detected with Benjamini-Hochberg corrected pairwise Wilcoxon Rank Sum tests between ‘inner petiole base’, ‘petiole’, and ‘peduncle’ samples (*P* ≤ 0.009). **B-D** – Images of plant samples bearing sporodochia. Orange triangles emphasize areas of sporodochial growth.

**Fig. 2.**
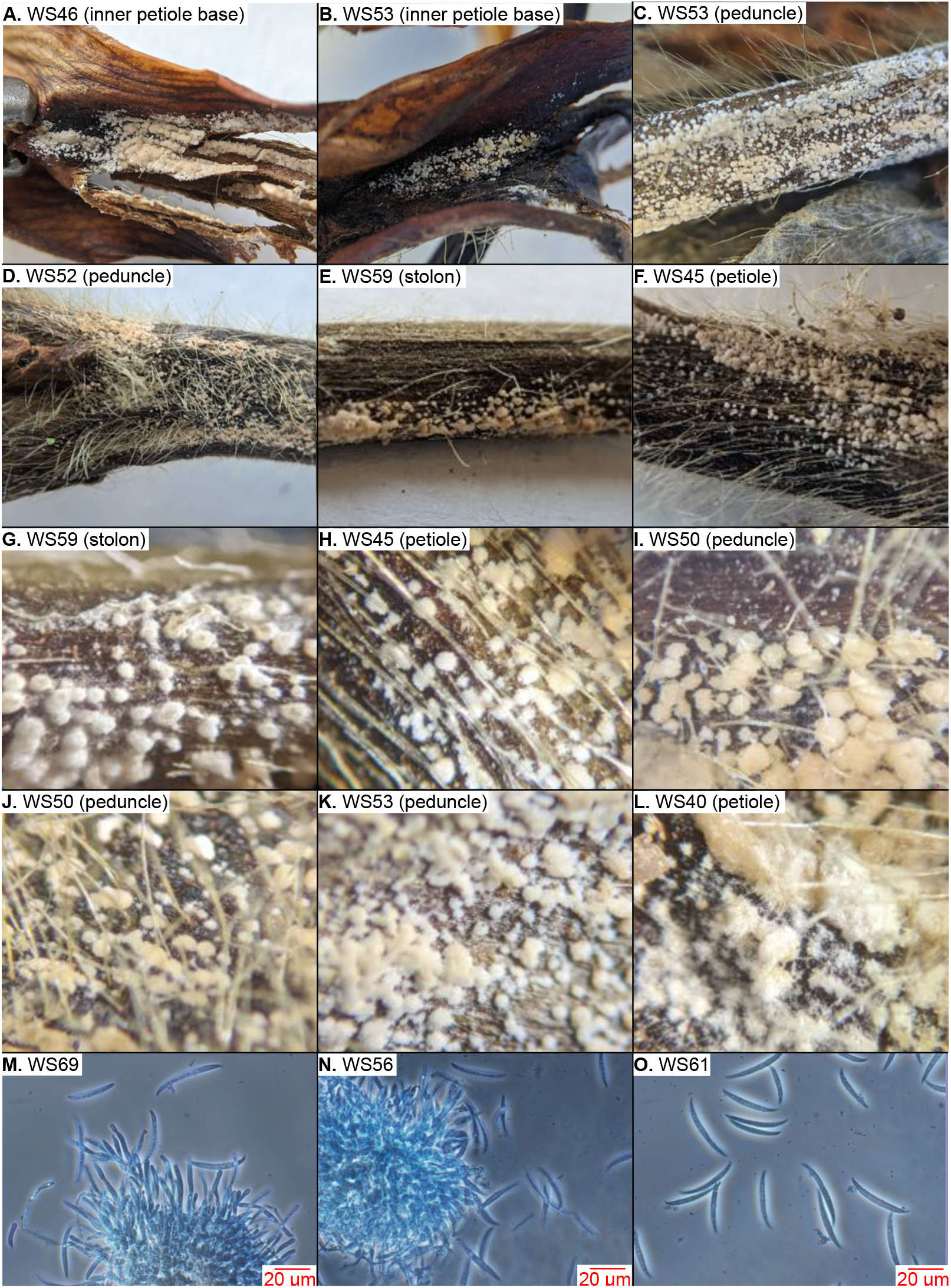
Images of sporodochia at 0×, 3×, and 400× magnification. **A-F** depict close-up photos of sporodochia taken without magnification. **G-L** show sporodochia at 3× magnification. **M-O** are images taken with a compound microscope at 400× magnification with phase contrast; macroconidia are visible in all three images and small sporodochia with conidiophores are also visible in M and N. The sample ID and tissue type are indicated in the upper left of each panel.

All four sporodochia samples were capable of shedding aerial conidia dispersed over short distances, albeit with varying efficiency (Figure 3). One sample only yielded one colony from the 2-hour wind treatment, but the others yielded 62, 56, and 99 colonies (Figure 3). Colonies recovered from vertically-oriented plates indicate longer-distance dispersal could be possible.

**Fig. 3.**
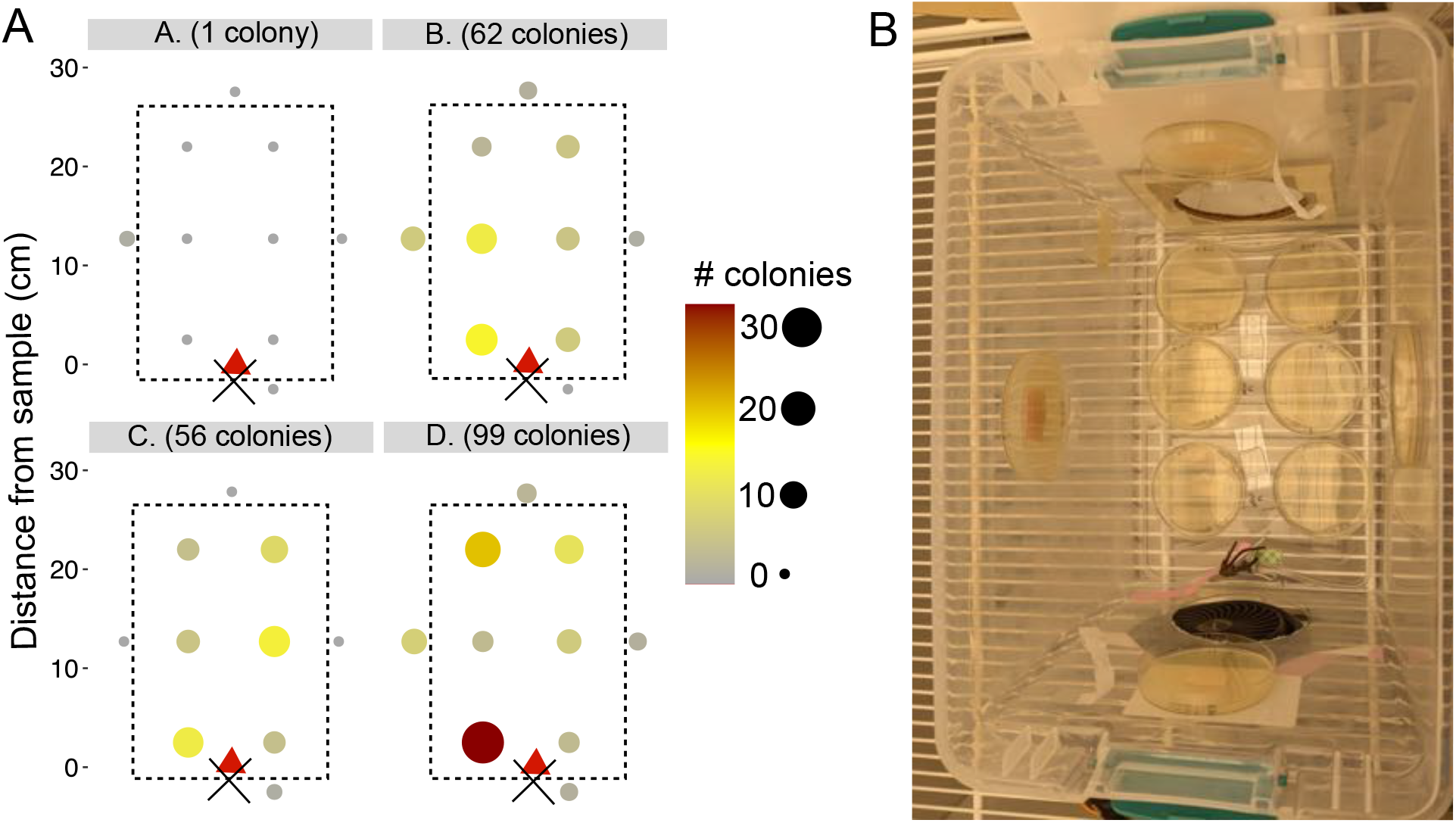
Aerial dispersal of macroconidia from sporodochia-bearing strawberry samples. **A** – The results of aerial dispersal for four samples are shown in sub-panels A-D. Dotted lines outline the bottom of the spore dispersal box. The position of the fan and sample are denoted with a black ‘X’ and red triangle, respectively. Circles depict the position of Petri plates and are color and size-coded according to the number of colonies they recovered. **B** – A photo of the spore dispersal box in the growth chamber.

These results demonstrated that sporodochia are commonly produced by *F. oxysporum* f. sp. *fragariae* on necrotic strawberry tissues during August and September in the Monterey Bay region of California. Sporodochia released airborne macroconidia that traveled short distances under moderate windspeed; travel over longer (>0.3 m) distances was not evaluated. Although it has long been understood that soilborne pathogens can spread on dust blown from one field to another, aerial dissemination of conidia must now be considered a significant risk for *F. oxysporum* f. sp. *fragariae*.

A major outstanding question is the efficiency of long-distance travel for these airborne conidia produced by *Fusarium oxysporum*: what is their range and do they pose risks for dissemination at regional, continental, or intercontinental scales? The efficiency of airborne long-distance travel would be primarily impacted by wind speeds at the ground level that dislodge spores, and the position of sporodochia on the plant (i.e. are they sheltered from wind by the canopy of the plant). These winds could be generated naturally by atmospheric conditions and augmented by the use of ‘bug vacuums’ that generate high windspeeds.

Many growers also use high-powered, tractor-mounted vacuums to remove *Lygus* spp. insects from strawberry plants. These vacuums are positioned just above the plants and can generate windspeeds between 7 and 54 km/hr in the plant canopy (Wells et al. 2020). Because *Lygus* spp. cause such extensive economic losses, the vacuums are commonly used on a weekly or twice-weekly basis for adequate control. These powerful windspeeds are more than sufficient to dislodge conidia and eject them at high speeds ∼6’ above ground level. There, conidia would be subject to further movement by natural atmospheric winds.

The maximum measured windspeed of 8.85 km per hour measured in our experiment is far less than normal afternoon windspeeds in the Monterey Bay around Salinas and Watsonville. In Salinas, afternoon windspeeds are often 21 km per hour, and windspeeds of 30 km per hour are not unusual. The windspeeds in Watsonville are typically less than in Salinas, but still often reach 14 km per hour during August and September. These windspeeds have sufficient force to dislodge conidia and could promote their transport over long distances.

While sporodochia on petiole bases are sheltered, we recorded sporodochia along petioles, peduncles, and stolons that would be more greatly influenced by air movement. Amazingly, the sporodochia on the single stolon sample covered 35 cm of length, a distance that could have extended beyond the plant canopy. Sporodochia lengths were highly variable on petioles and peduncles, but still regularly extended beyond 5 cm, a distance that remains within the canopy but accessible with high windspeeds. Also, necrotic tissues would bear dry wilted leaves that would be easily dislodged by mechanical disruption from weeding crews, bug vacuums, or pickers searching for fruit. Defoliation could further expose these sporodochia and promote dispersal.

An additional outstanding question regards the potential for arthropods to vector spores from one location to another. Although *F. oxysporum* f. sp. *cucumerinum* is not known to produce sporodochia, its conidia have been detected from air samples in infested greenhouses and on gnats and shore flies feeding at those locations (Scarlett et al. 2014). Mites and other arthropods have been implicated in the dispersal of *Fusarium* spp., but their contribution to dispersal has been difficult to disentangle from a role in promoting susceptibility to disease (Drakulic et al. 2017; Gamliel-Atinsky et al. 2009). It is unknown if arthropod species common in strawberry fields are attracted to sporodochia, and if so, what role they could play in long-distance dispersal.

The management implications of this discovery are profound and may explain previous observations of severe Fusarium wilt outbreaks in fields that were pre-plant broadcast fumigated with maximum rates. After broadcast fumigation, there is a period of a few days to several weeks where bare soil is exposed and could receive airborne conidia; this time period ends when the beds are fully formed and covered with plastic. Microbial abundance is decreased immediately after fumigation (Castellano-Hinojosa et al. 2022), and successfully germinating *F. oxysporum* f. sp. *fragariae* conidia may rapidly colonize the soil, as has been observed for *F. oxysporum* f. sp. *radicis-lycopersici* in steam pasteurized soils (Rowe et al. 1977). Sporodochia production and fumigation occur at the same time of year; mid-August until the end of September is peak fumigation season, and all our field collections were made during this time. Fields being prepared for planting in proximity to Fusarium wilt-afflicted fields may be better managed by drip fumigation which does not have a period of exposed, fumigated soil open to the air.

Successful management of any plant pathogen relies on an accurate understanding of its lifecycle and mechanisms for dispersal. Our observation of sporodochia produced by *F. oxysporum* f. sp. *fragariae* substantially improves our knowledge of the basic biology of this pathogen and its potential for spread. It also adds to a small but impactful body of literature on the potential for Fusarium wilt pathogens to be aerially dispersed. The fact that this was not documented in California for 16 years after the first observation suggests that there could be sporodochial phases for many other Fusarium wilt pathogens. Researchers of other *formae speciales* would benefit from close examination of diseased plants for these structures.

## Acknowledgements

We thank Jasmine Rodriguez for helping us to find diseased strawberry fields, and our funding sources: the California Strawberry Commission, the United States Department of Agriculture Agricultural Research Service, and National Institute of Food and Agriculture’s Specialty Crops Research Initiative award #: 2022-51181-38328.

## Literature Cited

Bilodeau, G. J., Koike, S. T., Uribe, P., and Martin, F. N. 2012. Development of an assay for rapid detection and quantification of Verticillium dahliae in soil. Phytopathology 102:331–343.

Burkhardt, A., Ramon, M. L., Smith, B., Koike, S. T., and Martin, F. 2018. Development of molecular markers to detect Macrophomina phaseolina from strawberry plants and soil. Phytopathology 108:1386–1394.

Burkhardt, A., Henry, P. M., Koike, S. T., Gordon, T. R., and Martin, F. 2019. Detection of Fusarium oxysporum f. sp. fragariae from infected strawberry plants. Plant Disease 103:1006–1013.

Castellano-Hinojosa, A., Boyd, N. S., and Strauss, S. L. 2022. Impact of fumigants on non-target soil microorganisms: a review. Journal of Hazardous Materials 427.

Drakulic, J., Bruce, T. J. A., and Ray, R. V. 2017. Direct and host-mediated interactions between Fusarium pathogens and herbivorous arthropods in cereals. Plant Pathology 66:3–13.

Gamliel-Atinsky, E., Freeman, S., Sztejnberg, A., Maymon, M., Ochoa, R., Belausov, E., and Palevsky, E. 2009. Interaction of the Mite Aceria mangiferae with Fusarium mangiferae, the Causal Agent of Mango Malformation Disease. Phytopathology 99:152–159.

Henry, P. M., Kirkpatrick, S. C., Islas, C. M., Pastrana, A. M., Yoshisato, J. A., Koike, S. T., Daugovish, O., and Gordon, T. R. 2017. The population of Fusarium oxysporum f. sp. fragariae, cause of Fusarium wilt of strawberry, in California. Plant Disease 101:550–556.

Henry, P. M., Pincot, D. D. A., Jenner, B. N., Borrero, C., Avilés, M., Nam, M. H., Epstein, L., Knapp, S., and Gordon, T. R. 2021. Horizontal chromosome transfer and independent evolution drive diversification in Fusarium oxysporum f. sp. fragariae. New Phytologist 230:327–340.

Katan, T., Schlevin, E., and Katan, J. 1997. Sporulation of Fusarium oxysporum f. sp. lycopersici on stem surfaces of tomato plants and aerial dissemination of inoculum. Phytopathology 87:712–719.

Koike, S. T., and Gordon, T. R. 2015. Management of Fusarium wilt of strawberry. Crop Protection 73:67–72.

Koike, S. T., Kirkpatrick, S. C., and Gordon, T. R. 2009. Fusarium wilt of strawberry caused by Fusarium oxysporum in California. Plant Disease 93:1077.

Komada, H. 1975. Development of a selective medium for quantitative isolation of Fusarium oxysporum from natural soils. Review of Plant Protection Research 8:114–125.

Miles, T. D., Martin, F. N., and Coffey, M. D. 2015. Development of rapid isothermal amplification assays for detection of Phytophthora spp. in plant tissue. Phytopathology 105:265–278.

Pincot, D. D. A., Poorten, T. J., Hardigan, M. A., Harshman, J. M., Acharya, C. B., Cole, G. S., Gordon, T. R., Stueven, M., Edger, P. P., and Knapp, S. 2018. Genome-wide association mapping uncovers Fw1, a dominant gene conferring resistance to Fusarium wilt in strawberry. G3: Genes, Genomes, Genetics 8:1817–1828.

R_Core_Team. 2013. R: A language and environment for statistical computing. R Foundation for Statistical Computing, Vienna, Austria.

Rekah, Y., Shtienberg, D., and Katan, J. 2000. Disease development following infection of tomato and basil foliage by airborne conidia of the soilborne pathogens Fusarium oxysporum f. sp. radicis-lycopersici and F. oxysporum f. sp. basilici. Phytopathology 90:1322–1329.

Rowe, R. C., Farley, J. D., and Coplin, D. L. 1977. Airborne spore dispersal and recolonization of steamed soil by Fusarium oxysporum in tomato greenhouses. Phytopathology 67:1513–1517.

Scarlett, K., Tesoriero, L., Daniel, R., and Guest, D. 2014. Sciarid and shore flies as aerial vectors of Fusarium oxysporum f. sp. cucumerinum in greenhouse cucumbers. Journal of Applied Entomology 138:368–377.

Schweigkofler, W., O’Donnell, K., and Garbelotto, M. 2004. Detection and quantification of airborne conidia of Fusarium circinatum, the causal agent of pine pitch canker, from two California sites by using a real-time PCR approach combined with a simple spore trapping method. Applied and Environmental Microbiology 70:3512–3520.

Wells, J., Fink, C., Edsall, M., Olivier, D., and Lin, J. 2020. Prototype Lygus spp. vacuum provides improved pest management in California strawberries. International Journal of Fruit Science 20:1019–1028.

